# Complex trait–environment relationships underlie the structure of Wisconsin forest plant communities

**DOI:** 10.1101/2020.11.17.387050

**Authors:** Andres G. Rolhauser, Donald M. Waller, Caroline M. Tucker

## Abstract

Plant species shift in abundance as environmental conditions change because traits adapt species to particular conditions. As a result, trait values shift along environmental gradients—the so-called trait–environment relationships. These relationships are often assessed by regressing community-weighted mean (CWM) traits on environmental gradients. Such regressions (CWMr) assume that local communities exhibit centered optimum trait–abundance relationships and that traits are not independent from one another. However, the shape of trait–abundance relationships can vary widely along environmental gradients—reflecting the interaction between traits and gradients—and traits are usually interrelated. Accounting for these complexities should improve our ability to accurately describe trait–environment relationships. We tested these ideas by analyzing how abundances of 185 herbaceous understory species distributed among 189 forested sites in Wisconsin, USA, varied in response to four functional traits (vegetative height-VH, leaf size-LS, leaf mass per area-LMA, and leaf carbon content) and six soil and climate variables. A generalized linear mixed model (GLMM) allowed us to assess how the shape of trait–abundance relationships changed along environmental gradients for the 24 trait–environment combinations simultaneously. We then compared the resulting trait–environment relationships to those estimated via CWMr. The GLMM identified five significant trait–environment relationships that together explained ∼40% of variation in species abundances across sites. Temperature played important roles with warmer and more seasonal sites favoring taller plants. Soil texture and temperature seasonality affected LS and LMA more modestly; these seasonality effects declined at more seasonal sites. Only some traits under certain conditions showed centered optimum trait– abundance relationships. Concomitantly, CWMr identified 17 significant trait–environment relationships including effects of temperature, precipitation, and soil on LMA as often reported in other studies. Despite this overidentification, CWMr failed to detect significant temperature-seasonality effects found in the GLMM. Modeling the complexity of how traits and environments interact to affect plant abundance allows us to identify and rank key trait– environment relationships. Although the GLMM model was more complex compared to single CWM regressions, it identified a simpler hierarchy of trait–environment relationships that accurately and reliably predicted responses of forest understory species to gradients in environmental conditions.

## Introduction

Understanding how plant species and communities respond to environmental gradients helps us predict future responses to global change (Kearney, Wintle, & Porter, 2010; Scheiter, Langan, & Higgins, 2013; Thuiller et al., 2008; van Bodegom, Douma, & Verheijen, 2014). In general, we expect plant phenotypes to be distributed in ways that reflect how species are adapted to local environments providing a mechanistic link between environmental change and community responses (Keddy, 1992; Lavorel & Garnier, 2002; Shipley, 2010; Violle et al., 2007). This approach has led to considerable empirical research seeking to characterizing how phenotypic traits relate to environmental gradients, i.e. so-called trait–environment relationships (Funk et al., 2017). However, reported trait–environment relationships vary widely in strength and sign (see reviews in Funk et al., 2017; Garnier, Navas, & Grigulis, 2016). This highlights the need to improve our understanding of these relationships if we are to make useful predictions.

The analytical tools we use to represent how traits vary along environment gradients must be suitably structured and complete enough to adequately capture the complexity that underlies ecological relationships. Mechanistically, trait–environment relationships arise because individual fitness or performance depend on how functional traits interact with environmental conditions (Laughlin & Messier, 2015; Laughlin, Strahan, Adler, & Moore, 2018; Shipley et al., 2016). In general, we say that traits and environments are related if the relative performance of species with different trait values change along an environmental gradient. This two-way interaction can be viewed from two perspectives (see also ter Braak, 2019). The site-centric perspective views how rules for local community assembly (Keddy, 1992), as represented by trait–performance relationships (Loranger, Munoz, Shipley, & Violle, 2018; Rolhauser & Pucheta, 2017), change as environmental conditions change (Fig. 1). The species-centric perspective instead views how traits modulate species’ abundance responses to environmental conditions, or environment–performance relationships (Fig. 1; Vesk, 2013). A corollary is that trait–environment relationships are inherently three-dimensional and cannot be readily reduced to, or inferred from, relationships viewed along individual trait or environmental axes. Rather, we must characterize how trait values and environmental conditions in combination affect performance if we are to reliably infer trait–environment relationships. A first step here is to acknowledge that these relationships can take different shapes.

**Figure 1.**
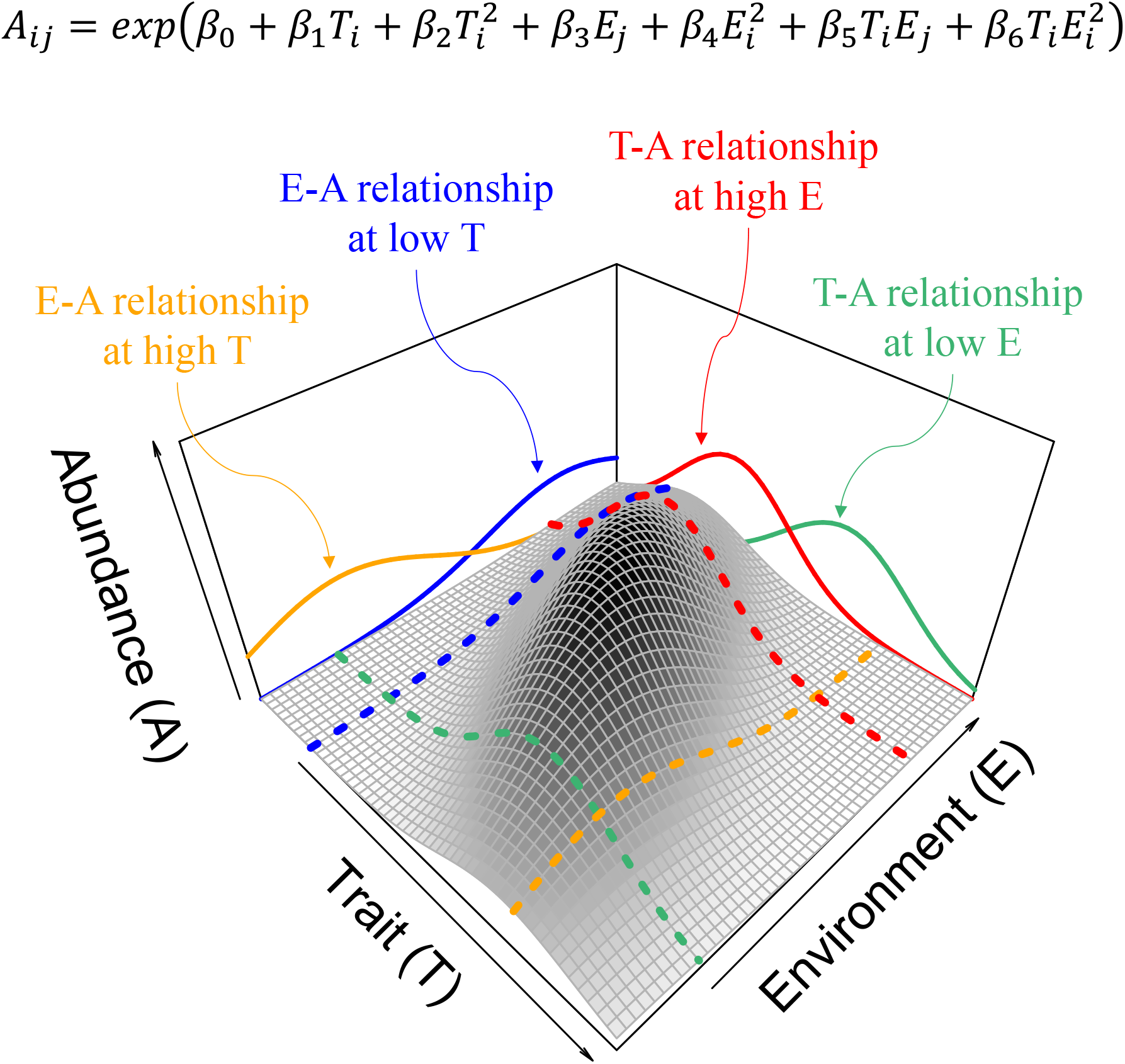
Fixed-effect structure of our GLMM approach modelling variation in the abundance of species *i* at site *j* (equation for *A*_*ij*_at the top). Here we used abundance as a proxy of performance. *A*_*ij*_(white to black surface) is modeled as an exponential function (through a log-link) of trait T and environmental variable E. Parameters *β*_1_ and *β*_2_ are the linear and quadratic effects of T on A; *β*_3_ and *β*_4_ are the linear and quadratic effects of E on A; and *β*_5_ and *β*_6_ are the linear and quadratic effects of E on T (see Statistical methods). Combining the log link with negative quadratic terms (with *β*_2_ and *β*_4_ set to −1 and −0.5, respectively, here), this model reproduces unimodal trait–abundance (orange and blue) and environment–abundance (green and red) relationships. In this example, the result is a directional and negative T-E relationship (*β*_5_ and *β*_6_ set to −1 and 0, respectively). Dotted lines are modeled relationships at different T or E levels while continuous lines are their counterparts projected onto 2-D planes.

If species are best adapted to particular environmental conditions, environment– performance relationships should be humped or unimodal reflecting intermediate optima (Austin, 1999; Curtis, 1959; Ter Braak & Prentice, 1988; Whittaker, 1967). Environment–performance relationships can also be bimodal, among other reasons, because of intense negative interspecific interactions (Austin, 1999; Minchin, 1987; Mueller-Dombois & Ellenberg, 1974). Similarly, we expect unimodal trait–performance relationships when functional trade-offs lead to optimum trait values among coexisting species (Muscarella & Uriarte, 2016; Rolhauser, Nordenstahl, Aguiar, & Pucheta, 2019; Rolhauser & Pucheta, 2017). Trait–performance relationships can be bimodal (or multimodal) when within-site environmental heterogeneity favors the functional divergence of competitive, dominant species (Rolhauser et al., 2019; Rolhauser & Pucheta, 2017).

Directional relationships emerge as particular cases when performance maxima (in optimum relationships) or minima (in bimodal relationships) occur at the extremes or outside the range of observed trait and environmental values (Rolhauser et al., 2019; Ter Braak & Prentice, 1988). Unlike trait–performance and environment–performance relationships, theoretical discussions on the shape of trait–environment relationships are rare. Both conceptual (Garnier et al., 2016; Laughlin & Messier, 2015; Shipley et al., 2016) and empirical studies (Moles et al., 2014, 2009; e.g. Niinemets, 2001; Wright et al., 2017, 2004) largely assume directional relationships. Nonetheless, some empirical work exists to show that trait–environment relationships can be nonlinear (Bello et al., 2013; Laughlin, Fule, Huffman, Crouse, & Laliberté, 2011).

Studies often simplify the three-dimensional problem to infer trait–environment relationships. Those that assess how the performance of individual species respond to environmental conditions remain rare and typically assume directional trait–performance and environment–performance relationships (e.g. Laughlin et al., 2018). Direct regressions of trait values against environmental variables (trait–environment regressions) make the stronger assumption that the abundance or presence of a species at a given site indicates their performance (Funk et al., 2017; Garnier et al., 2016). In many cases, community-weighted mean (CWM) traits are used as the response variable, referred to here as CWM regressions or CWMr (Miller, Damschen, & Ives, 2019), but there are also regressions of site-specific species trait averages (e.g. Dong et al., 2020) and individual trait values (Laughlin, Joshi, van Bodegom, Bastow, & Fulé, 2012).

Trait–environment regressions have two further conceptual limitations. First, because they do not explicitly model abundance, they cannot account for possible variations in trait– abundance and environment–abundance relationships. For example, CWM values represent the adaptive value of a trait only at sites showing centered optimum trait–abundance relationships (Muscarella & Uriarte, 2016) despite this being a special case (Rolhauser & Pucheta, 2017). Second, they tend to evaluate traits one at a time using separate regressions, implicitly assuming independence between traits and their responses to environments. However, traits are inherently interrelated within integrated phenotypes (Murren, 2012). Evolutionary biologists have long recognized the importance of using multivariate approaches when trying to tease apart the adaptive significance of correlated traits (Lande & Arnold, 1983). Both complexities reduce the power of trait–environment regressions to reliably detect important relationships, potentially generating false-negative conclusions. In addition, CWMr are known to inflate type I errors by falsely flagging too many trait–environment associations as significant (Miller et al., 2019; Peres-Neto, Dray, & ter Braak, 2017).

We evaluated trait–environment relationships across forest understory communities in Wisconsin, USA, using a generalized linear mixed model (GLMM) approach (ter Braak, 2019; Vesk, 2013; Warton et al., 2015). This approach can fully account for and incorporate the underlying complexities discussed above. Compared to CWMr, GLMM better balances type I error control and power allowing it to have lower rates of both false positives and false negatives (Miller et al., 2019; ter Braak, 2019). Our model allows all three functional relationships (trait– abundance, environment–abundance, and trait–environment) to be nonlinear (Figs. 1, S3). The model also allows us to evaluate multiple traits and trait–environment relationships simultaneously. In doing so, this approach seeks to explain species-abundance variation across sites on the basis of traits, environmental gradients and their interactions (Fig. 1). We capitalize here on the existence of three large and interrelated datasets. The vegetation data matrix includes detailed data on the abundances of 185 species distributed across 189 sites (Waller, Amatangelo, Johnson, & Rogers, 2012). The functional trait data include values for ten traits characterizing tissues (e.g. leaf mass per area, or LMA), organs (e.g. leaf size, or LS), and whole plants (e.g. maximum vegetative height, or VH). The environmental data include 14 variables describing both abiotic (e.g. mean annual temperature and soil sand content) and biotic (tree basal area) environmental conditions. Functional traits like VH, LS, and LMA are often studied for their effects on plant performance and show wide variation along abiotic and biotic gradients (see review in Pérez-Harguindeguy et al., 2013). Finally, we compare the trait–environment relationships resulting from the single GLMM to those found using CWM regressions. We find that the GLMM not only explains a large proportion of the variability in species abundances but also that it improves and simplifies the process of inferring the most important trait–environment relationships.

## Material and Methods

### Study area

Wisconsin covers a large area (169,639 km^2^) but displays little variation in elevation (range: 177-595 m). Its north temperate climate is highly seasonal. It has conspicuous north-south gradients in summer and winter temperatures and east-west gradients in seasonality and rainfall (reflecting the moderating maritime influence of Lake Michigan to the East). The vegetation is dominated by deciduous forests that can be divided into two floristic provinces with the prairie-forest province to the southwest and northern hardwoods to the north. These are separated by a marked “tension zone” running from the SE to the NW edges of the state (Curtis, 1959; see Fig. S1 in Supporting Information).

### Vegetation sampling

Data were collected by resurveying many forest stands (sites) first surveyed by J.T. Curtis and his students in the 1950s (see Fig. S1 in Supporting information). Only sites that remained less disturbed (i.e. with an intact forest canopy, no nearby edges, and few signs of understory disturbance) were resurveyed. Resurveys occurred between 2000 and 2012 using Curtis-era and updated methods covering similar areas (Waller et al., 2012; see Appendix S1 for details). Briefly, all vascular plants were tallied within each of many replicate (20-403) 1-m^2^ quadrats to estimate species abundances (frequency) at each site. In total, 536 species were found at 293 sites. We restricted our attention to the commonest 185 species for which we had access to locally collected trait data. We then selected sites for analysis with high trait coverage (>80% of all species occurrences within quadrats). We discarded a few sites missing full environmental data, yielding a total of 189 sites.

### Environmental variables

Descriptions of the four climatic variables, ten soil variables and one biological variable (tree basal area) appear in Appendix S1. Climate variables for each site derive from 10-year averages for the period 1995-2004. Replicate soil samples from each site were combined and analyzed for cations and physical properties. We estimated tree cover (basal area) at each site using tree surveys. We used pairwise correlations and principal components analyses to select relatively independent subsets of climatic and soil variables and so avoid collinearity among predictors (correlation coefficients between all retained predictors were below 0.5, Appendix S1). This generated six predictors: mean annual temperature (MAT), mean annual precipitation (MAP), temperature seasonality (i.e. the standard deviation of mean monthly temperatures, abbreviated TSD), soil nitrogen content (%N), soil sand content (%Sand), and tree basal area (BA). Before analysis, environmental variables were transformed as needed to reduce the weight of extreme values, then standardized to zero mean and unit variance (Appendix S1).

### Plant traits

Amatangelo *et al*. (2014) characterized functional trait variation in the common herbaceous species in these communities using standard methods (see Appendix S1 for trait descriptions and methods). Briefly, at least 12 individuals (4+ plants from each of 3+ sites) were collected in Wisconsin between 2008 and 2014 and processed following standardized protocols (Pérez-Harguindeguy et al., 2013). From ten traits available, we selected four for analysis with low pair-wise correlations (<0.3; Table S3). These are: vegetative height (VH), leaf size (LS, calculated as the product of length and width), leaf mass per area (LMA), and leaf carbon content (LCC). Traits were transformed as needed to reduce the weight of extreme values and standardized (Appendix S1).

### Statistical methods

#### GLMM

We sought to infer the shape and significance of trait–environment relationships while modeling meaningful site and species responses to, respectively, trait and environmental axes (Fig. 1). We used a negative binomial generalized linear mixed model (GLMM) to model the abundance of species *i* at site *j* (*A*_ij_) as a function of the selected traits and environmental variables using the natural logarithm link function. *A*_ij_ is a count variable calculated as the frequency of quadrats in which species *i* was present at site *j*. Negative binomial models directly estimate an overdispersion parameter from the data (Agresti, 2015). Before describing the full multivariate model, we outline our analysis for one trait and one environmental variable (both standardized). Given the log link function, the fixed effects in the GLMM are (cf. Fig. 1):

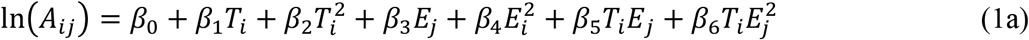

where *β*_0_ is the *y*-intercept of the fitted surface. Since trait *T* is centered at zero, *β*_1_ and *β*_2_ respectively estimate the mean slope (or “gradient”) and the mean curvature of the trait– abundance relationship (see e.g. Aiken, West, & Reno, 1991) for the particular case when *E*_*j*_is set to its average value (zero). That is, *β*_1_ measures the direction (positive or negative) and strength of trait *T*_i_’s effect on species abundance. Negative values of *β*_2_ indicate “n” shaped (optimum or unimodal) relationships, while positive values indicate “u” shaped (bimodal) relationships. Similarly, *β*_3_ and *β*_4_ reflect the mean slope and curvature of the environment– abundance relationship (ter Braak, 2019). Due to the combination of quadratic effects (*β*_2_ and *β*_4_) and the log link associated with the negative binomial distribution, Equation (1a) can reproduce bell-shaped trait–abundance and environment–abundance relationships (Figs. 1, S3a). While doing this, we assess trait–environment relationships by estimating interactions between *T* and *E*. These parameters quantify how different environmental conditions modulate the mean effect of the trait on the response variable (Laughlin et al., 2018). In our model, these environmentally mediated trait effects are estimated by *β*_5_ and *β*_6_, as explained below.

Rearranging Eqn. 1a to gather terms for the mean (linear) effect of trait *T*_i_ on *A*_ij_yields:

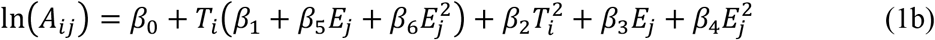

Then, the environment-dependent mean slope of trait–abundance relationships (denoted *φ*_*j*_) is:

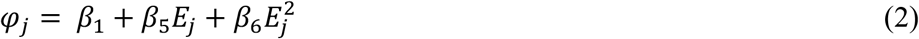

In this quadratic function, *β*_5_ and *β*_6_ are respectively the mean slope and mean curvature of the trait–environment relationship (Fig. S3). Indeed, since 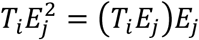, *β*_6_ estimates how the strength of the trait–environment interaction depends on values of the environment (see equation 1a), a quadratic relationship. Importantly, the properties of the fitted surface (e.g. the position of optimum trait values) change along the environmental gradient depending on *β*_5_ and *β*_6_ (Fig. S3).

The full GLMM included four traits (VH, LMA, LS, LCC) and six environmental variables (MAT, MAP, TSD, %Sand, %N and BA). Following Equation (1a), the fixed effects included linear and quadratic terms for all traits and environmental variables as well as all trait– environment interactions. We formulated random effects after ter Braak (2019) including random intercepts and slopes for all traits and environmental variables (i.e. MLM3 sensu ter Braak, 2019). The full model in R code notation is shown in Appendix S1. In this formulation, we estimate random effect slopes for each trait, *T*, for each site (denoted *b*_*(T)j*_here). We similarly estimate random effect slopes for each environmental variable, *E*, for each species (*c*_*(E)i*_) (ter Braak, 2019). We accounted for search-effort by including the log number of quadrats sampled at each site, used here as an offset (Kéry, 2010, pp. 188-189). We fitted the GLMM using the R-package glmmTMB (Brooks et al., 2017). We calculated marginal and conditional R^2^ (the proportion of the total variance explained by fixed effects and by both fixed and random effects, respectively) following the delta method (Nakagawa, Johnson, & Schielzeth, 2017). We implemented R^2^ calculations using the r.squaredGLMM function in the R-package MuMIn (Barton, 2019). We visualized uncertainty around estimates of *φ*_*j*_(for a given *T*-*E* combination) by plotting overall site effects calculated as the sum of the fixed (*φ*_*j*_) and the random slopes associated with the corresponding trait (*b*_*(T)j*_) (ter Braak, 2019).

#### Comparing GLMM and CWMr approaches

We calculated community-weighted means (CWM) as usual:

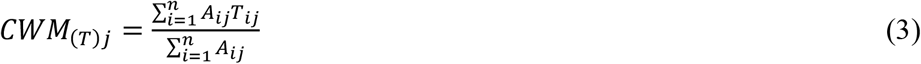

where *A*_ij_ and *T*_ij_ are the abundance and the standardized trait value for species *i* in site *j*. We then related CWM values to standardized environmental variables (*E*_*j*_) through univariate regressions (separate for each trait). Including both linear and quadratic terms for *E*_*j*_, leads to the general form:

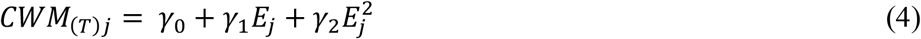

We fitted CWMr models using the least squares method implemented in the lm function in R.

To compare the GLMM and CWMr approaches, we compared the statistical significance of homologous model parameters for each trait–environment combination, i.e. *β*_5_ vs *γ*_1_ and *β*_6_ vs *γ*_2_ (cf. equations 2 and 4). We divided test statistics (*z* and *t* in the case of GLMM and CWMr, respectively) based on whether they led to the rejection of the null hypothesis or not and compared these between approaches (see Appendix S1). There are several possible outcomes for each trait–environment relationship tested (Table S4; Laughlin et al., 2018). When both CWMr and GLMM show non-significant effects, they agree that the trait has little adaptive value along that environmental gradient (Outcome #1). A non-significant trait–environment relationship with CWMr that is significant with GLMM suggests that the trait does have adaptive value along the gradient, but that CWMr lacked the sensitivity to detect it (Outcome #2). If, conversely, the CWMr trait–environment interaction is significant but the GLMM term is not, the CWMr result may be spurious or indirect as found by others who noted highly inflated type I error rates (Miller et al., 2019; Peres-Neto et al., 2017) (Outcome #3). Finally, if both methods identify a significant relationship with the same sign, we can conclude the trait has adaptive value along that gradient (Outcome #4). However, significant relationships for both approaches might also emerge with opposite signs (Outcome #5, Table S4).

## Results

### GLMM

#### Overview

The full model included 113 parameters (69 fixed effects, 43 random effects, and the overdispersion parameter) to explain abundance across 34,965 species–site combinations. The marginal R^2^ of this model was 0.385 with a conditional R^2^ of 0.977. We focus on trait– environment relationships, but full model results appear in Appendix S2. Quadratic terms between trait and environment effects on abundance were all negative, indicating unimodal curvilinear relationships (Table S5). Importantly, removing either all quadratic or all interaction terms from the full model increased the AIC >>2 (Table S6), supporting their retention in the final model.

#### Trait–environment relationships

The GLMM model identified five trait–environment relationships as significant, defined as those combinations of a trait and an environmental variable where either or both interaction parameters (*β*_5_, *β*_6_) were significant (*p*<0.01; Table S5). Three of these relationships were largely directional (only *β*_5_ was significant; blue lines in Fig. 2). The interaction between mean annual temperature (MAT) and vegetation height (VH) was the strongest (*β*_5_=1.031; SE=0.146). All remaining significant linear interactions (*β*_5_) had slopes below 0.34 (Table S5; note unitless standardized coefficients). Warmer sites (higher MAT) favored taller plants (Fig. 2, top line second plot, and Fig. 3, a1, a2). Height also increased linearly with temperature seasonality (TSD: *β*_5_=0.328, SE=0.082; Fig. 2, top line third plot, and Fig. 3, b1 and b2). In addition, leaf size (LS) declined linearly as soil sand content increased (*β*_5_=-0.266, SE=0.075; Fig. 2, row 2 column 4, and Fig. 3, c1, c2).

**Figure 2.**
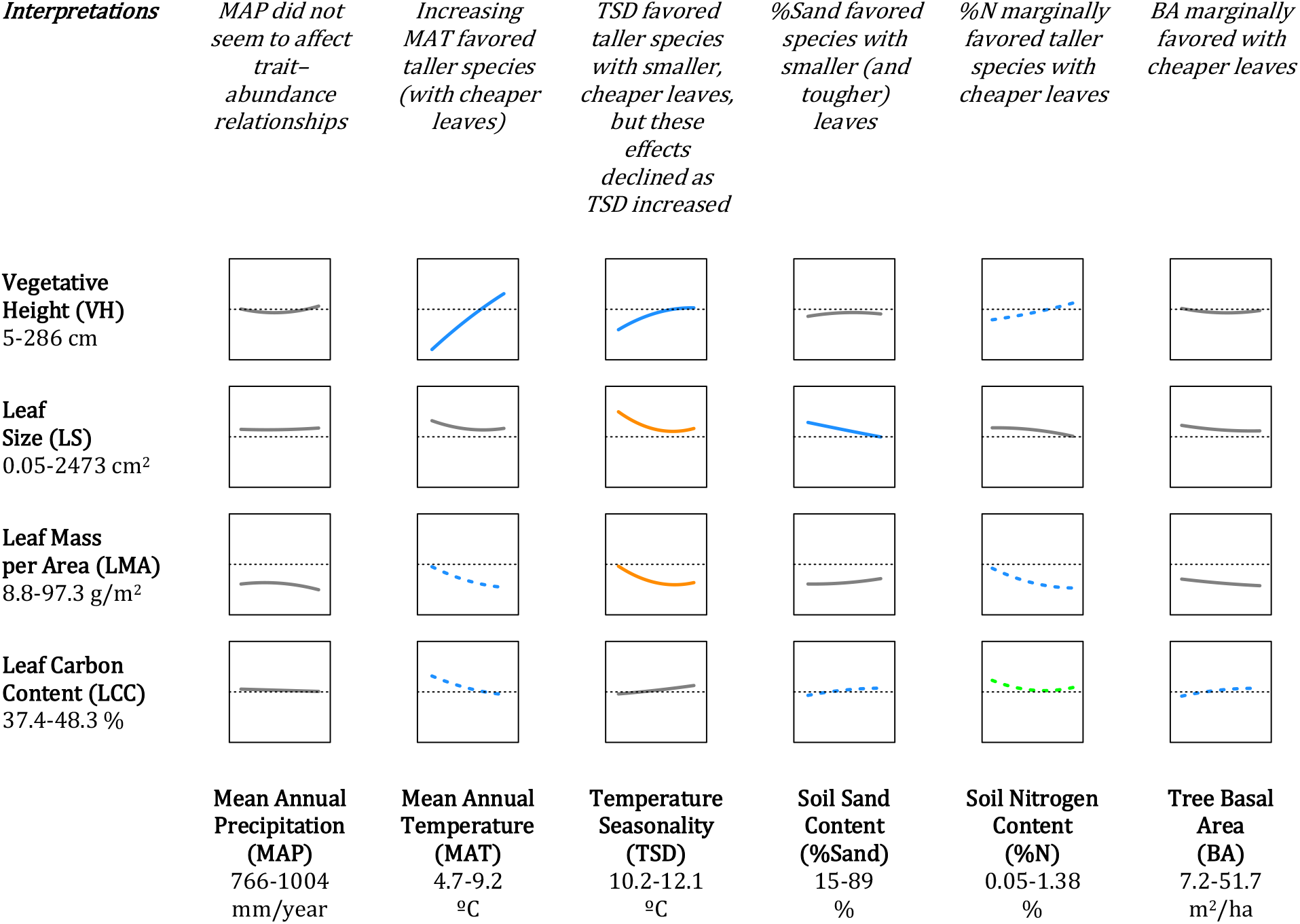
Trait–environment relationships as estimated by the GLMM (Equation 2). Relationships between traits (y-axes) and environmental variables (x-axes) are plotted based on estimates of their the linear (*β*_5_) and quadratic (*β*_6_) coefficients (Table S5). The observed range of traits and environmental variables are shown below the axis labels. Blue lines show relationships where only linear effects are significant (i.e. *β*_5_≠0), green lines show those where only nonlinear effects are significant (i.e. *β*_6_≠0), and orange lines those where both effects are significant. Dashed lines show relationships of marginal significance (0.01<*p*<0.05) while grey lines show nonsignificant ones. The dotted horizontal lines indicate y=0. “Interpretations” above describe overall patterns (with marginally significant patterns parenthesized).

**Figure 3.**
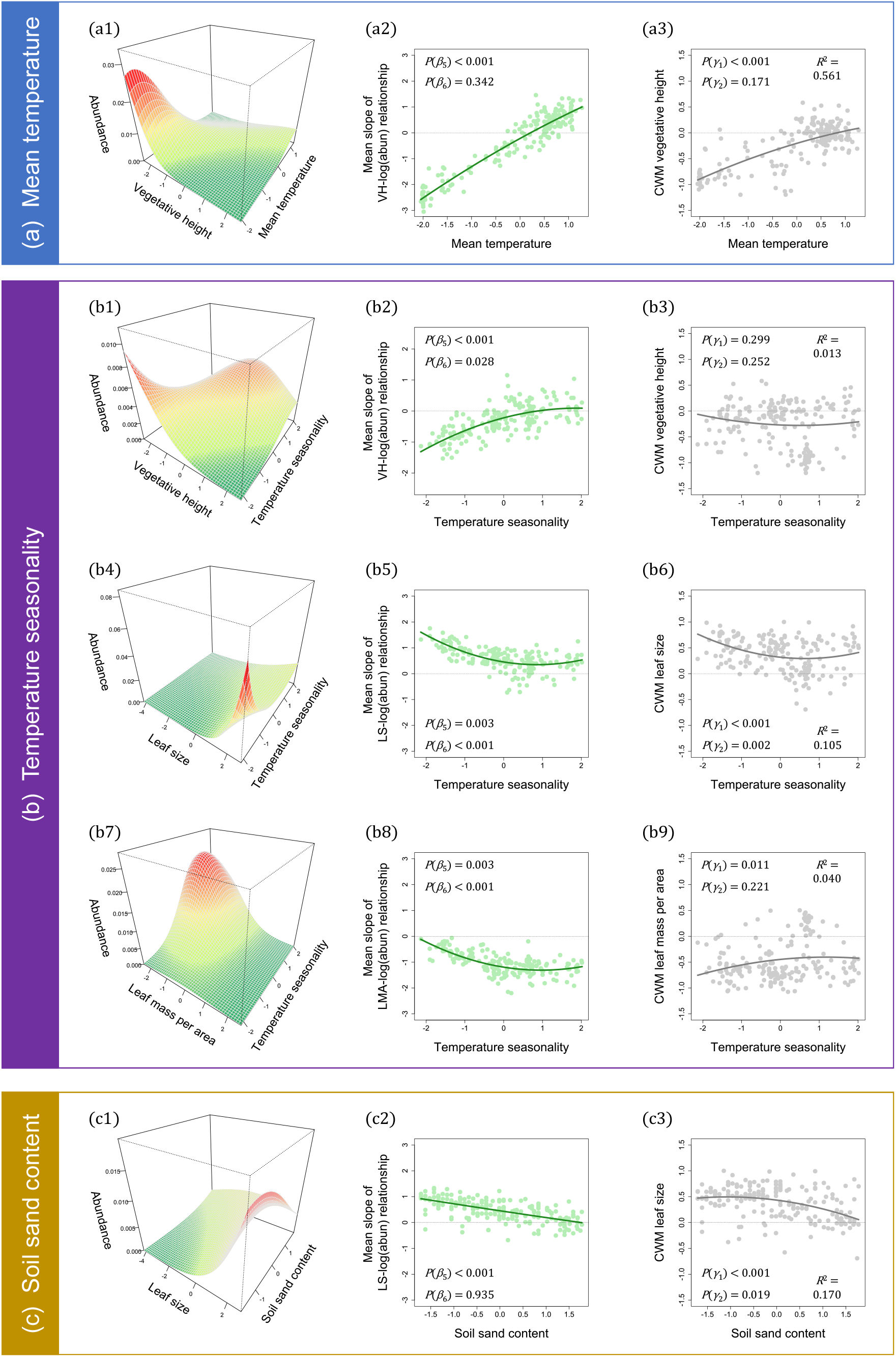
Details of significant interactions (*p*<0.01) from the GLMM visualized for (a) temperature seasonality, (b) mean annual temperature, and (c) soil sand content. Surfaces in the first column show species abundance fitted as a function of trait values and environmental condition (holding other variables constant at their mean value). Plots in the second column show average slopes (*B* in Equation 2) of the given trait–abundance relationship (green line). *B* depends on *β*_5_ and *β*_6_ which estimate the linear and quadratic components of the trait– environment relationships, respectively. Each green dot represents the overall effect of site *j*, i.e. the sum of the fixed (*φ*_*j*_) and the random effect slope associated with a trait (*b*_*(T)j*_). Plots in the third column show traditional community-weighted mean regressions (CWMr’s) between observed CWM trait values (grey dots) and observed environmental variables (Equation 3; dark grey line fitted by Equation 4). Significance levels are shown for *γ*_1_ and *γ*_2_ (the linear and quadratic terms) and CWMr overall R^2^. The dotted horizontal lines indicate y=0. Surfaces are colored to enhance the perception of relief.

Sites with more seasonal temperatures (TSD) supported species with smaller leaves (LS, *β*_5_=-0.242, SE=0.082) and lower leaf mass per area (LMA, *β*_5_=-0.239, SE=0.082; Fig. 3: b4, b5, b7, and b8). In both cases, these decreases were nonlinear (*β*_6_≠0, orange lines, Fig. 2, column 3; Table S5) with decelerating effects of seasonality on LS and LMA as seasonality increased.

Other linear trait–environment interactions were marginally significant (0.01<*p*<0.05, blue dashed lines in Fig. 2). These included increases in plant height with soil N (row 1, column 5), declines in LMA and LCC with increasing temperatures (rows 3 and 4, col 2), and increases in LCC as soil sand (row 4, column 4) and basal area increased (row 4, column 6). LCC also showed a marginally significant nonlinear response to soil N (green dashed line, row 4, column 5). Surprisingly, annual precipitation (MAP) did not appear to affect any of these traits (column 1).

Note that significant trait–environment interactions mean that the shapes of trait– abundance relationships change along environmental gradients (Fig. 3, column 1). We did detect roughly centered optimum trait–abundance relationships (the form assumed by the CWMr approach) for some trait–environment combinations, sometimes for restricted regions along environmental gradients. These centered optimum patterns appeared most notably for VH around the mean MAT (Fig. 3a1) and at mid-to-high TSD (Fig. 3b1). These relationships, however, become directional away from these environmental regions. We found largely directional or flat trait–abundance relationships for the remining three significant trait–environment relationships (Fig. 3: b4, b7 and c1).

### Comparing GLMM and CWMr approaches

In contrast to the five significant trait–environment relationships (of 24) identified using GLMM, the CWMr approach identified 17 regressions where either or both parameters (*γ*_1_ and *γ*_2_) showed *p*<0.01 (Table S7). Twelve of these were only directional (only *γ*_1_≠0; e.g. LMA–MAT), four were curved with a directional component (both *γ*_1_≠0, *γ*_2_≠0; e.g. LMA–%N), and one was curved but lacked directionality (only *γ*_2_≠0; LCC–MAP; Table S7).

Test statistics for linear trait–environment parameters from both the GLMM and CWMr approaches (*β*_5_ and *γ*_1_) paralleled each other (R=0.61; *t*=3.572, df=22, *p*=0.002; Fig. 4a). Test statistics for the quadratic effects (*β*_6_ and *γ*_2_) were less similar (R=0.41; *t*=2.095, df=22, *p*=0.048; Fig. 4b). Both approaches agreed that six linear effects (including all four interactions with BA, Fig. 4a) and 16 quadratic effects lacked significance (Fig. 4b), representing Outcome #1. In addition, three linear effects of environmental conditions on plant traits (LS–TSD, LS–%Sand, and VH–MAT) and one quadratic term (LS–TSD^2^) emerged as significant in both approaches (Outcome #4). Reassuringly, we found no cases where the two approaches yielded opposite conclusions (Outcome #5). However, inferences often differed conspicuously between GLMM and CWMr. Two linear and one quadratic effect emerged as significant in the GLMM but not in the CWMr approach (Outcome #2). Among the five significant trait–environment relationships in the GLMM (second column, Fig. 3), two were largely missed by CWMr. Both involved temperature seasonality (third column, Fig. 3). More worrying, many apparent Type I errors occurred as 13 linear and four quadratic terms judged significant by the CWMr approach lacked significance under the GLMM (Outcome #3).

**Figure 4.**
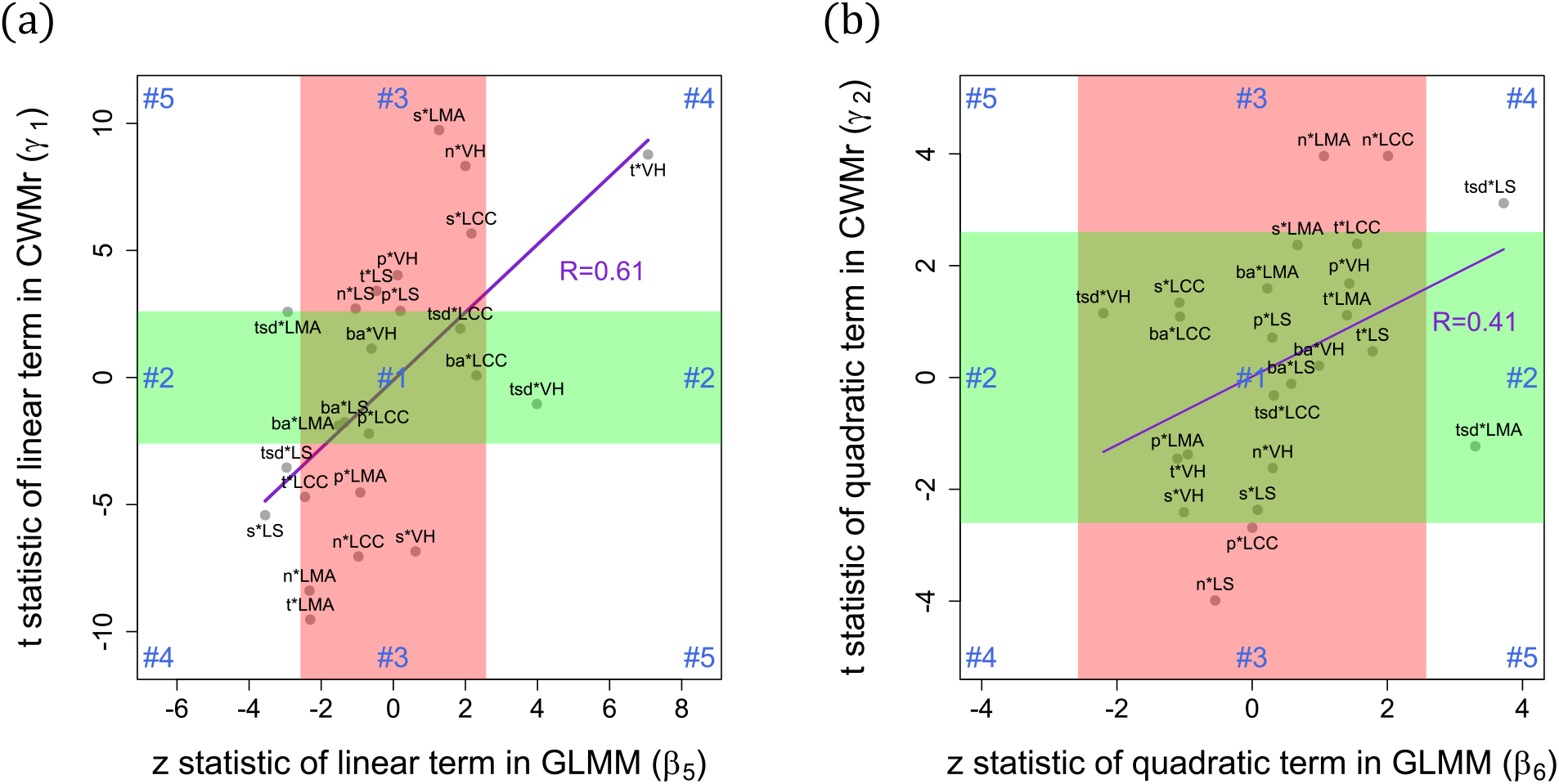
Scatterplots relating test statistics from univariate CWM regressions to statistics for homologous terms from the GLMM. Plot (a) compares linear terms (*γ*_1_ and *β*_5_) while plot (b) compares quadratic terms (*γ*_2_ and *β*_6_). Each data point corresponds to one of the 24 trait– environment relationships. The purple lines are fitted to all 24 points. Corresponding Pearson’s correlations (R) show that estimated linear terms agree better between the two approaches than the quadratic terms. Green and red rectangles show ranges of test statistic values failing to reject the null hypothesis (of no significant effect) under the CWM and GLMM models, respectively. The five Outcome codes shown in blue refer to those outlined in Material and Methods and Table S4. Environmental variable and trait codes as in Fig. 2, except that p = mean annual precipitation, t = mean annual temperature, s = soil sand content, n = soil nitrogen content.

## Discussion

Trait–environment relationships provide a useful tool to understand and predict community responses to environmental change, assuming their accuracy is not compromised by using an inappropriate modeling approach. Unlike simpler community-weighted mean regressions (CWMr), generalized linear mixed models (GLMM) are more flexible and comprehensive in being able to incorporate complex and sometimes interacting trait–environment relationships. In these understory herbaceous communities, the CWMr approach overidentified many relationships as significant that lacked significance in the more functionally complete and informed GLMM model. Just five relationships emerged as significant (of the 24 evaluated) in the GLMM, including two missed by the CWMr approach. These, however, were sufficient to explain nearly 40% of the total variation in abundance among 185 plant species distributed over 189 sites. Recent GLMM applications achieved similar explanatory power (between 0.25 and 0.49) in analyzing presence-absence data (Löbel, Mair, Lönnell, Schröder, & Snäll, 2018; Pollock et al., 2018). Our work confirms that mixed-effects models provide powerful tools to explain not only distributions of species but also their abundances across large spatial scales.

### Prominent trait–environment relationships

The interaction between mean annual temperature (MAT) and vegetative height (VH) emerged as the strongest trait–environment relationship in the GLMM in terms of the size and significance of its slope (*β*_5_). It was also the only relationship where the slope of the trait– abundance relationship (*φ*_*j*_) switched signs along the environmental gradient (Fig. 3), reflecting clear shifts in the rank order of species abundances. The positive height–temperature relationship matches global patterns (Moles et al., 2014, 2009). It may be explained by the fact that warmer climates favor taller plants as they compete for light (Falster & Westoby, 2003; Givnish, 1982) and/or that shorter plants are less prone to freeze-embolism in cold climates (Moles et al., 2009).

Temperature seasonality tended to increase both vegetative height and plant abundance, confirming how many understory species thrive in highly seasonal environments. This may reflect how Spring and Fall periods of higher light in these deciduous forests and warmer forest floor conditions favor understory plant growth. If the growth of many species is favored at more seasonal sites, these sites could generate the more competitive, crowded conditions that favor taller plant species (Falster & Westoby, 2003; Givnish, 1982).

Both leaf size (LS) and leaf mass per area (LMA, the inverse of specific leaf area) declined as temperature seasonality (TSD) increased. Low-LMA leaves are “cheaper” in terms of dry-mass investment and therefore tend to be fast-growing, although short-lived (Poorter, Niinemets, Poorter, Wright, & Villar, 2009). Faster-growing species with low-LMA leaves might thus be favored at sites with short Spring windows of full light, e.g., shorter seasons (Kikuzawa, Onoda, Wright, & Reich, 2013; Mason & Donovan, 2015). Given that leaves often start to generate carbohydrates at about half their final size (Turgeon, 1989), shorter seasons might also favor smaller leaves for their ability to start exporting carbohydrates sooner than larger leaves.

Leaf size (LS) declined at sites with sandier soils in agreement with global patterns where water-limited environments tend to favor plants with smaller leaves (Wright et al., 2017). The standardized coefficient (*β*_5_) for the LS–%Sand interaction was only a quarter the size of that for the VH–MAT interaction. Thus, soil properties may play a smaller role in forest plant community assembly than climate factors at our scale of observation.

Among environmental factors, only temperature seasonality (TSD) interacted nonlinearly with plant traits with effects of TSD on LS and LMA declining as seasonality increased. Nonlinear patterns also emerged in other studies (Bello et al., 2013; Laughlin et al., 2011), suggesting that trait–environment relationships may not be constant enough along some environmental gradients to expect consistent patterns to emerge. Such nonlinear effects could help to explain some of the inconsistencies found in the literature in terms of the strength and sign of these relationships (Funk et al., 2017; Garnier et al., 2016). The scale of observation and extent of sampling, in particular, could affect outcomes if studies differ in which portions of environmental gradients they sample (Pollock et al., 2018).

### Missing trait–environment relationships

Many regional to global-scale studies find that levels of precipitation can strongly affect plant traits (Moles et al., 2014, 2009; Niinemets, 2001; Wright et al., 2017). In contrast, we found no trait–precipitation interactions. This may reflect the fact that these Wisconsin sites span only 5% of the 0 to 4500 mm/year range spanned at the global scale. In contrast, soil sand varied from 15% to 89% among sites, allowing it to affect leaf size (and leaf carbon content somewhat). Thus, sand content may better reflect water availability for plants among our sites.

Basal area, our surrogate for understory light, only affected LCC weakly, perhaps reflecting how widely understory light levels vary within sites (Canham, Finzi, Pacala, & Burbank, 1994). In contrast, basal area emerged as the single best predictor of species occurrences among sites (Beck et al., 2020). The weak linear effects we found for soil N (favoring taller plant species with cheaper leaves) could also reflect high local variability in soil conditions or perhaps that soil N mainly affects plants through interactions with other environmental variables. Such environment–environment interactions were omitted in our already complex GLMM.

Relationships between temperatures and leaf characteristics have attracted the attention of many ecologists (e.g. the “leaf economics spectrum”, Wright et al., 2004). The negative LMA– MAT relationship we found was marginally significant, consistent with other regional-scale studies finding weakly negative or non-significant relationships (Balazs et al., 2020; Dong et al., 2020; Laughlin et al., 2012, 2018; Mason & Donovan, 2015; Rosbakh, Römermann, & Poschlod, 2015). Such weak effects are surprising given the strong negative effects found under controlled growth conditions (Poorter et al., 2009). LMA responded more to temperature seasonality in the GLMM. In larger, especially global, datasets, mean temperature and seasonality are so closely correlated (see e.g. Kikuzawa et al., 2013) that we cannot decouple their separate effects.However, in our dataset, the MAT–TSD correlation was mild (−0.37; Table S2), suggesting that the weak MAT effect on LMA was not an artifact of seasonality soaking up variation in the model. Much remains to be understood regarding the individual and interactive effects of MAT and seasonality on LMA (Kikuzawa et al., 2013).

### GLMM vs. CWMr approaches

The large number of significant but potentially spurious relationships identified here using the CWMr approach supports conclusions based on simulated data that CWMr inflates Type I errors (Miller et al., 2019; Peres-Neto et al., 2017). Many terms in the CWMr identified as significant (including LMA with MAP, MAT, sand, and %N, LS with MAP and MAT, and VH with MAT and %N) lacked significance in the more nuanced and complete GLMM model, reflecting Outcome #3. These relationships have been commonly reported in the literature, suggesting some reports may reflect spurious false positives. In the case of LMA, a widely studied trait, the lack of significance in the GLMM cannot be ascribed to collinearity with competing traits as all inter-trait correlations were small (<0.18, Table S3). It is more likely to reflect how important it is to account for complex interrelationships among multiple predictor variables.

Conversely, two trait–environment relationships featuring temperature seasonality emerged as highly significant in the GLMM (with VH and LMA) but were not significant in the CWMr (Outcome #2). These hidden relationships emerged in a model that flexibly handled trait– abundance and environment–abundance relationships while controlling for effects of other trait– environment relationships. These results suggest that CWMr approaches are not only prone to false positives but also to false negatives. This might account for why such TSD relationships are under-reported in the literature, particularly in cases were MAT and TSD are highly correlated.These differences in inference are particularly striking given that both models used exactly the same data.

### Strengths, limitations, and future directions

The GLMM approach provided deeper and more reliable insights than could be attained using separate CWMr, which led to both apparent false positive and false negative conclusions. Modeling trait–environment combinations together in one model accounts for non-independent effects on species abundance arising from collinearity. Including quadratic effects incorporates more realistic shapes for functional relationships. The nonlinear trait–environment interactions we introduced in our GLMM approach emerged as particularly important for characterizing effects of temperature seasonality. Paradoxically, the more complex GLMM identified a simpler hierarchy of relationships relative to the CWMr approach with temperature effects strongly driving variation in plant height while soil texture (and temperature seasonality) affected leaf traits more modestly.

We interpreted *φ* _*j*_ as reflecting how strongly trait values are “favored”, “selected” or “filtered” along particular environmental gradients (Loranger et al., 2018; Rolhauser & Pucheta, 2017; Shipley, 2010). We implicitly assumed that trait–environment effects on plant abundance act independently of other processes like dispersal and demographic stochasticity. Nevertheless, we acknowledge that stochasticity can play important roles in community dynamics (Alonso, Etienne, & McKane, 2006; Hubbell, 2001) and likely contributes to the remaining 60% of the variation in species abundances that remained unexplained in our GLMM. Dispersal limitation clearly limits the presence or abundance of some species in the fragmented forests of southern Wisconsin (Rogers, Rooney, Hawbaker, Radeloff, & Waller, 2009). However, the traits we used here are more related to local plant performance (see Pérez-Harguindeguy et al., 2013) than to dispersal among forest patches. For example, differences in plant height clearly contribute to both competitive success and dispersal, but mostly at fine scales (Gómez, 2007).

Some have criticized analyzing trait–environment relationships from distributions of plants rather than individual or population-level performance (e.g. demographic rates) as phenomenological (Laughlin & Messier, 2015; Laughlin et al., 2018). This criticism may be misplaced if problems associated with phenomenological approaches arise rather by collapsing or aggregating the abundance dimension using CWMr or other trait–environment regressions. It would therefore be useful to compare GLMMs based on abundance to studies of individual or population-level performance to assess how well they match. Mismatches between these approaches could reflect key links between performance and abundance, a keystone of community ecology that remains only loosely supported (Adler, Fajardo, Kleinhesselink, & Kraft, 2013; McGill, Enquist, Weiher, & Westoby, 2006; Shipley et al., 2016).

## Acknowledgements

The vegetation surveys and studies of trait variation were supported by NSF grants DEB 023633, DEB 0717315, and a Dimensions of Biodiversity award (DEB-1046355). We also thank J. Beck for providing comments on an early version of the manuscript and Jeremy Ash for his assistance in the compilation of climatic data. The authors declare no competing interests.

## Author contributions

AGR and CMT conceived the ideas and designed the study, with input from DMW, who also provided the data. AGR analyzed the data and led the writing. All authors contributed critically to the drafts and gave final approval for publication.

## Data availability

Should the manuscript be accepted, the data supporting our results will be archived in Figshare and the data DOI will be included at the end of the article.

## Supporting Information

**Appendix S1** – Supplementary Material and Methods

**Appendix S2** – Supplementary Results

