## Supplementary information for "Complex trait–environment relationships underlie the structure of Wisconsin forest plant communities"

### Appendix S1 – Supplementary Material and Methods

#### *Study area*

The sampled forest stands are located in Wisconsin, USA (Fig. S1). These stands vary in composition and history, but all were originally identified and sampled by Prof. J.T. Curtis and his students in the 1950s and described in his classic Vegetation of Wisconsin book (Curtis 1959). That work focused on describing and categories the state's plant communities as a vehicle to examining species and community responses to gradients in environmental conditions, paralleling our goal here. It also provides details on the physical geography and glacial history of Wisconsin, soils, and climatic conditions across the state. Briefly, the state covers a broad area (169,639 km$^2$) but only a limited range of elevation (177-595 m). It was mostly glaciated during the most recent (Wisconsin) glaciation except for the "Driftless area" to the southwest. Its highly seasonal north temperate climate shows a conspicuous north-south gradient in summer and winter temperatures and an east-west gradient in seasonality (reflecting the moderating maritime influence of the Great Lake Michigan to the East). It spans two floristic provinces: the prairie-forest province to the southwest and the northern hardwoods province to the north, separated by marked "tension zone" running from the SE to the NW edges of the state (Curtis 1959; Fig. S1). More than 182 species spanning many orders reach either the southern or northern limit of their distribution here reflecting abrupt changes in climatic conditions across this tension zone.

**Figure S1**. Distribution of the 189 sites in Wisconsin, USA (red points, panel a), where the two floristic provinces (prairie-forest and northern hardwoods) and the intermediate "tension zone" are also shown. The size of symbols represents mean annual temperature (MAT) in panel b and temperature seasonality (TSD) in panel c.

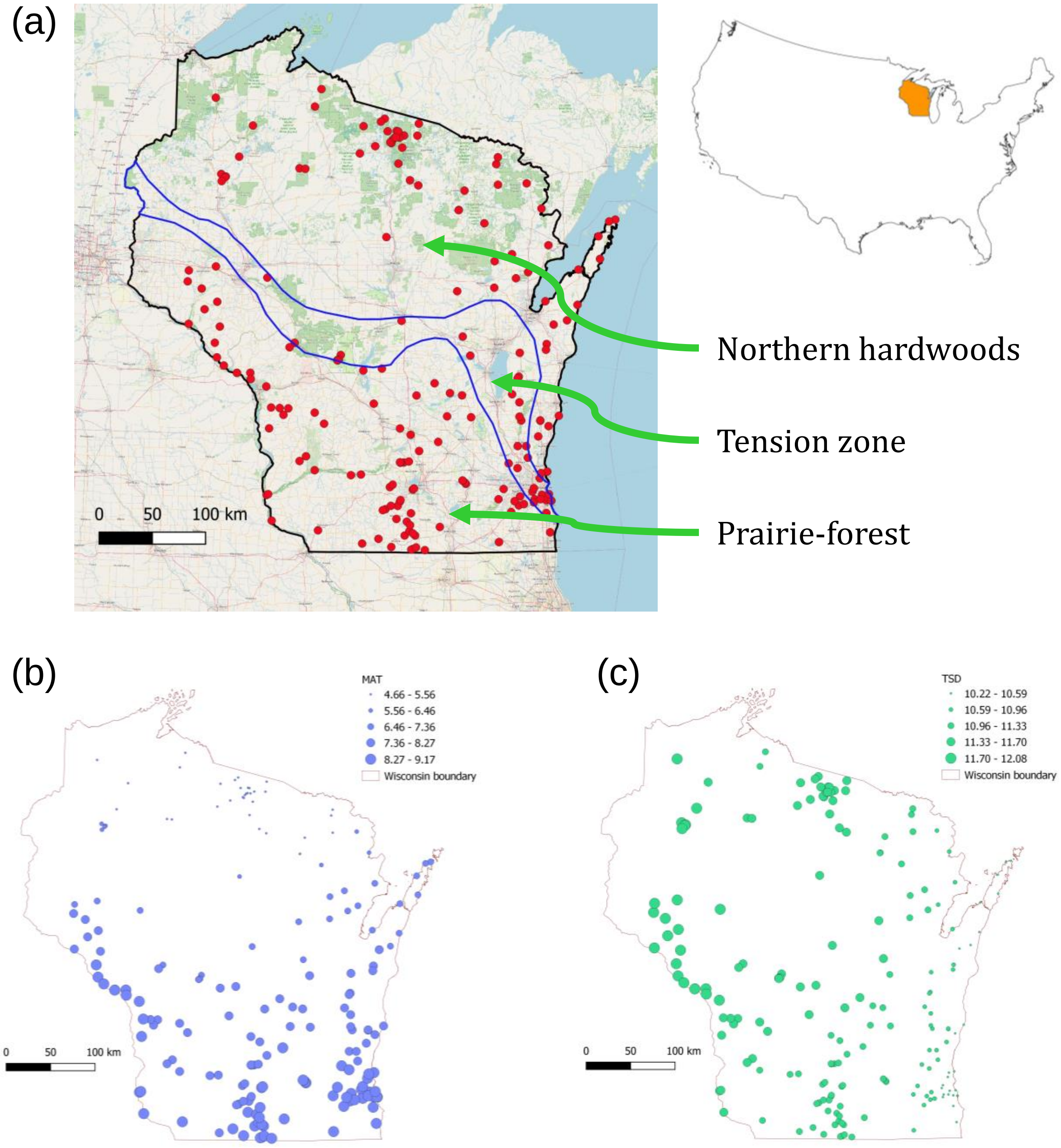

#### *Vegetation Sampling*

Our data derive from contemporary surveys of 276 forest stands originally surveyed by J.T. Curtis and his students in the 1950s (Fig. S1). These 2000s era resurveys were designed to measure and characterize the nature and extent of changes in plant community composition and diversity in northern upland forests (NUF) (Rooney et al. 2004; Wiegmann and Waller 2006), southern upland forests (SUF) (Rogers et al. 2008, 2009), southern lowland forests (SLF) (Johnson and Waller 2013; Johnson et al. 2014, 2016), and pine barrens forests (PB) (Li and Waller 2015, 2016, 2017). These resurveys used both Curtis-type original methods and extensions of these to cover more quadrats (Waller et al. 2012; Table S1). Only sites that remained relatively undisturbed (i.e. with an intact forest canopy, no nearby edges, and few signs of understory disturbance) were resurveyed.

**Table S1**. Summary of forest plant communities surveyed in the 2000s. PEL refers to the original Wisconsin Plant Ecology Laboratory methods (Cottam and Curtis 1949). "Area covered" is a conservative estimate based on the areas of the transects plus a 20m buffer on all sides.

| Forest type | Survey years | Number of sites | Mean number of quadrats / site | Survey method | Area covered (ha) |
|---|---|---|---|---|---|
| Beech (B) | 2003-2004 | 20 | 86 | Original PEL | 2.6 - 4.0 |
| Northern Upland (NUF) | 2000-2004 | 49 | 97.6 | Strip transects & original PEL | 0.8 – 4.0 |
| Pine Barrens (PB) | 2012 | 5 | 50 | Parallel transects | 1.1 |
| Southern Lowland (SLF) | 2007-2008 | 31 | 39 | Parallel transects | 1.4 |
| Southern Upland (SUF) | 2002-2005 | 84 | 104.2 | Original PEL | 2.6 - 4.0 |
| | Totals: | 189 | 16,715 | | |

These resurveys characterized understory composition by tallying the frequency at which each species occurred across 42 to 120 1-m$^2$ quadrats. The number and spatial distribution of quadrats were chosen to account for the species diversity present at each site (Johnson et al. 2008). These were arranged regularly either along two or three large U's (matching original 1950s methods), along parallel transects covering a similar area, or along six 20-m strips of adjacent quadrats (Table S1). Resurveys of the SUF used matching PEL protocols except that two or three large squares or U's spaced 30-40 m apart were used to increase sample sizes (Rogers et al. 2008). The NUF-B sites were resurveyed using 20 adjacent 1-m$^2$ quadrats arranged as six 20-m long strips (Rooney et al. 2004). The 40 southern lowland forest sites are located along 13 major rivers and streams primarily in southern Wisconsin (Fig. S1). These were selected to be low, flat, and poorly drained with no signs of recent fire, cutting or grazing (Johnson and Waller 2013). Resurveys of these sites arranged 42 1-m$^2$ quadrats spaced every 7.5 m along six parallel 50-m transects located 20 m apart. Resurveys of the Pine Barrens sites located in Wisconsin's central sand plains (Fig. S1) surveyed 50 1-m$^2$ quadrats arranged with a spacing of 5 m along five parallel 50-m transects spaced 20 m apart (Li and Waller 2015). The areas sampled clearly affect the number of species detected, but the exact layout and spacing of quadrats have little effect on estimates of diversity in these forests (Johnson et al. 2008).

#### *Environmental variables*

We compiled data on four climatic variables relevant to large-scale vegetation dynamics (Prentice 1986), i.e. mean annual temperature (MAT), mean annual precipitation (MAP), temperature seasonality (TSD), precipitation seasonality (PCV). Raw precipitation and temperature data were provided by the PRISM Climate Group (Daly et al. 2004). We extracted values for each site and averaged across the 10-year period between 1995 and 2004. Soils were characterized using pooled 300cc samples from the top 10 cm of soil (A horizon) taken from 3-10 dispersed points at each site. Soil samples were then analyzed by the Wisconsin state Soil and Plant Analysis Laboratory for soil texture (% sand, silt, and clay) and constituents (organic matter, pH, N, P, Ca, Mg, and K).

Environmental variables were transformed when necessary to reduce the weight of extreme values and then standardized to zero mean and unit variance. Namely, soil nitrogen content (%N) was log transformed while basal area (BA) was square root transformed. Correlations were below 0.4 for all climatic pairs except for mean annual temperature (MAT) and precipitation seasonality, i.e. the coefficient of variation of monthly precipitation (Table S2). Principal components analysis of the soil variables returned a two-dimensional solution explaining ~70% of the variance (Fig. S2). The first principal component (PC1) mostly reflects fertility (pH, % organic matter, percentage of N, and parts per million of Ca and Mg) and explains 50% of the variance. PC2 mostly reflects soil texture (% sand vs. silt) and explains 19% of the variance. We chose %N and %Sand to represent variation in soil conditions among these sites. Therefore, our final list of environmental variables (which will be used as predictors in our models) included: mean annual temperature (MAT), mean annual precipitation (MAP), temperature seasonality (i.e. the standard deviation of mean monthly temperatures, abbreviated TSD), soil nitrogen content (%N), soil sand content (%Sand) and tree basal area (BA). None of the pairwise correlations between these predictors were considered problematic (Table S2), i.e. they were all bellow 0.5 (see e.g. Dormann et al. 2013). MAT and TSD were highly correlated with spatial gradients: MAT declined with latitude (increasing southward) while TSD declined with longitude (increasing westward, Table S2).

**Table S2.** Pearson correlations between climate, soil variables and spatial gradients. Correlations >0.5 are shown in orange. The variables in green were chosen to enter the model as fixed-effect, explanatory variables.

| | Lat | Long | sqrt(BA) | MAT | MAP | TSD | PCV | %Sand | ln(%N) |
|---|---|---|---|---|---|---|---|---|---|
| Latitude | 1 | | | | | | | | |
| Longitude | -0.198 | 1 | | | | | | | |
| Basal Area [sqrt(BA)] | -0.207 | 0.097 | 1 | | | | | | |
| Mean annual temperature (MAT) | -0.940 | 0.127 | 0.197 | 1 | | | | | |
| Mean annual precipitation (MAP) | -0.468 | -0.305 | 0.068 | 0.336 | 1 | | | | |
| Temperature seasonality (TSD) | 0.434 | -0.926 | -0.124 | -0.366 | 0.119 | 1 | | | |
| Precipitation seasonality (PCV) | -0.678 | -0.358 | 0.147 | 0.725 | 0.331 | 0.181 | 1 | | |
| Soil sand content (%Sand) | 0.475 | 0.062 | 0.068 | -0.475 | -0.275 | 0.083 | -0.315 | 1 | |
| Soil nitrogen content [ln(%N)] | -0.351 | 0.226 | 0.028 | 0.421 | -0.063 | -0.332 | 0.170 | -0.423 | 1 |

**Figure S2**. Principal components analysis of soil data. There is a strong gradient of fertility (~56%) of variation), and a second one (~14%) related to texture.

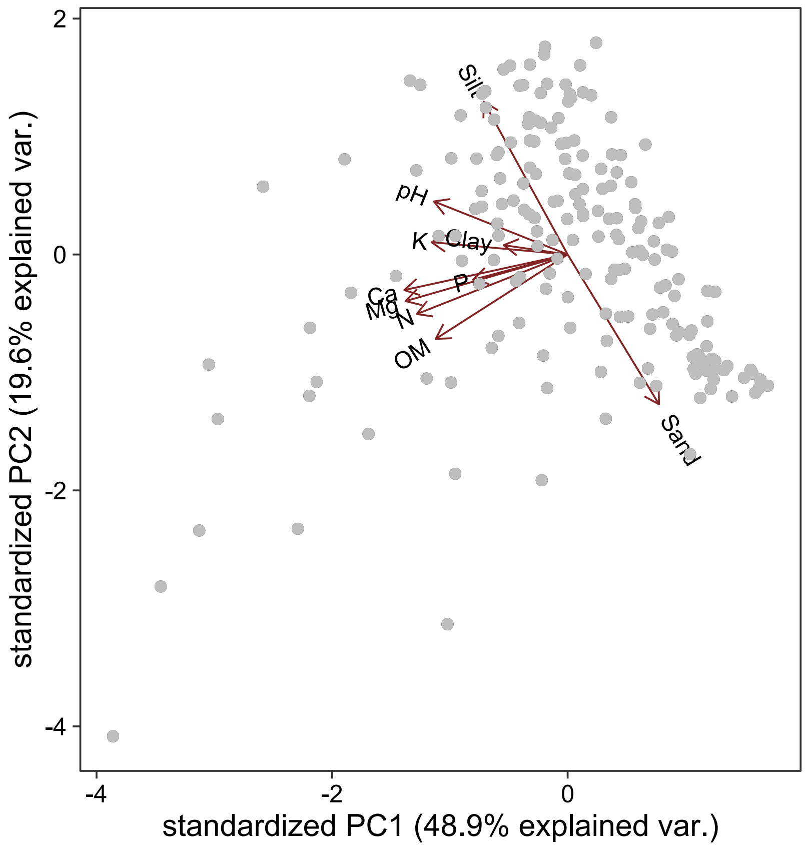

#### ***Plant traits***

The traits data derive from plants collected in Wisconsin and processed in Waller's lab at the University of Wisconsin between 2008 and 2014 as initially described by Amatangelo et al. (2014). We measured traits on at least 12 individuals (four individuals from each of three sites) following standardized protocols (Pérez-Harguindeguy et al. 2013). Leaf height was measured vertically from the ground to maximum (unstretched) leaf height. For leaf traits, we collected two young but fully expanded leaves from each individual (including both basal and cauline leaves for species where these were markedly different). We measured lamina leaf thickness using an Ames Pocket Thickness Gauge (B.C. Ames, Framingham, MA USA), avoiding major veins. We then removed petioles and scanned the leaves on a flatbed scanner. These images were analyzed for area, length, and width using Image J (Schneider et al. 2012). The leaves were then weighed, dried for at least 48 hours at 60$^{o}$ C, and reweighed to determine leaf mass per area ratio (LMA, i.e. the inverse of specific leaf area) and leaf dry matter content (LDMC). We averaged trait values across all individuals, providing one value per species. Although some within-species variation exists for most traits, this is small relative to among-species variation (D. Waller, unpublished data). We pooled samples of dried leaf material for each species and analyzed them for nitrogen content on a Flash EA Analyzer (ThermoScientific, Waltham, Massachusetts, USA). For carbon chemistry and extended nutrient analyses, we sent leaf material pooled across sites to the Wisconsin state Plant and Soil Analysis Lab to obtain one value per species for leaf P, K, Ca, Mg, ash, and carbon content.

Plant traits were transformed when necessary to reduce the weight of extreme values and then standardized to zero mean and unit variance. Namely, LMA, maximum vegetative height (VH), leaf size (LS, the product of leaf length and width), leaf length (LL) and leaf width (LW), leaf thickness (LT), leaf carbon:nitrogen ratio (CN) were log transformed. Leaf nitrogen content (LNC) and leaf dry matter content (LDMC) were logit transformed, while LCC was kept untransformed. LMA was highly correlated (i.e. correlation >0.5) LT, leaf CN, LNC, and LDMC (Table S3). LS was highly correlated with LL and LW, while VH and LCC showed relatively low correlations (<0.4) with other traits (Table S3). Our final list of traits used as predictors in our models include: VH, LS, LMA and LCC. All pairwise correlations between these predictors were below 0.3 (Table S3).

**Table S3.** Pearson correlations between ten functional traits measured in 185 herbaceous species. Correlations >0.5 are shown in orange. In green we show the traits (and their correlations) that were chosen to enter the model as fixed-effect, explanatory variables.

| | ln(VH) | ln(LS) | ln(LL) | ln(LW) | ln(LMA) | ln(LT) | ln(CN) | logit(LNC) | logit(LDMC) | LCC |
|---|---|---|---|---|---|---|---|---|---|---|
| ln(VH) | 1 | | | | | | | | | |
| ln(LS) | 0.274 | 1 | | | | | | | | |
| ln(LL) | 0.392 | 0.647 | 1 | | | | | | | |
| ln(LW) | 0.107 | 0.879 | 0.206 | 1 | | | | | | |
| ln(LMA) | 0.175 | -0.175 | 0.043 | -0.252 | 1 | | | | | |
| ln(LT) | -0.043 | 0.005 | 0.035 | -0.016 | 0.620 | 1 | | | | |
| ln(CN) | -0.266 | -0.334 | -0.146 | -0.337 | 0.627 | 0.327 | 1 | | | |
| logit(LNC) | 0.275 | 0.368 | 0.164 | 0.371 | -0.646 | -0.377 | -0.987 | 1 | | |
| logit(LDMC) | 0.297 | -0.224 | 0.063 | -0.327 | 0.598 | -0.087 | 0.559 | -0.528 | 1 | |
| LCC | 0.005 | 0.155 | 0.083 | 0.147 | -0.005 | -0.245 | 0.257 | -0.100 | 0.283 | 1 |

#### *Nonlinear trait–environment relationships assessed by the GLMM*

According to our model (equation 1, main text; see also Fig. S3), the properties of the fitted surface, such as positions of optimum trait values, change along the environmental gradient depending on $\beta_5$ and $\beta_6$. For instance, if $E$ affects $T$ positively ($\beta_5 > 0$) and linearly ($\beta_6 = 0$), the position of the optimum on the trait axis will change at constant pace towards higher trait values as $E$ increases (Figs. S3a, b). This linear trajectory of the optimum trait is reflected by the linear relationship between $\varphi_j$ and $E_j$ (Fig. S3c). When $\beta_6 \neq 0$ (e.g. negative, leading to a n-shaped relationship) and $\beta_5>0$, the optimum's position increases at a decreasing rate (Fig. S3d, e), and this curved trajectory of the optimum trait is reflected by the nonlinear relationship between $\varphi_j$ and $E_j$ (Fig. S3e).

**Figure S3.** Hypothetical surfaces of species abundance as a function of a trait and an environmental variable for linear (a) and nonlinear (d) trait-environment relationships holding al other variables constant at their average value. Panels (b) and (e) are contour plots of surfaces in panels (a) and (d), respectively. The white arrows were added manually to show that the ridge between the two peaks in (b) is rectilinear, while that in panel (d) is curved. The average slope of the trait-abundance relationship (denoted $\varphi_j$) captures these linear (c) and nonlinear (f) trait-environment relationships. The equations that define each panel are place at the top of the figure. Parameters defining the surface in panel (a) were $\beta_2 = -1$ and $\beta_5 = 1$, and the remining were set to zero. In panel (d), parameters were $\beta_1 = 1.3$ , $\beta_2 = -1$, $\beta_5 = 1$, and $\beta_6 = -0.3$ (the value of $\beta_1$was chosen to produce a two-peak surface similar to that in the linear scenario).

$$Abundance_{ij} = exp(\beta_0 + \beta_1 T_i + \beta_2 T_i^2 + \beta_3 E_j + \beta_4 E_i^2 + \beta_5 T_i E_j + \beta_6 T_i E_i^2) \qquad \varphi_j = \beta_1 + \beta_5 E_j + \beta_6 E_j^2$$

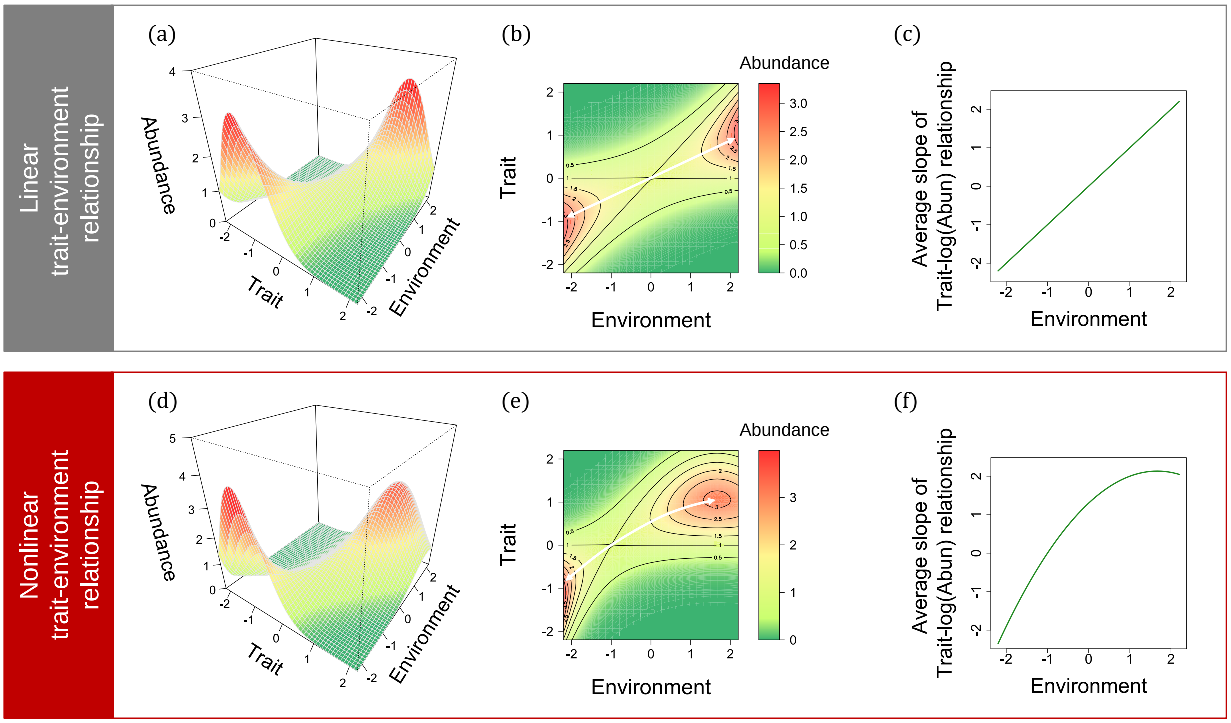

#### *Full GLMM in R code notation*

The full GLMM included four traits (VH, LMA, LS, LCC) and six environmental variables (MAT, MAP, TSD, %Sand, %N and BA) and their interactions (Following equation 1a) as fixed effects. We included random intercepts and slopes for all traits and environmental variables (ter Braak 2019). We also included the log number of quadrats sampled per site as an offset to account for search-effort heterogeneity (Kéry 2010, pp188-189). We fitted this GLMM using the R-package glmmTMB (Brooks *et al.* 2017). The model in R code notation is:

```
glmmTMB(abundance~(MAP + MAT + TSD + %Sand + %N + BA + I(MAP^2)
+ I(MAT^2) + I(TSD^2) + I(%Sand^2) + I(%N^2) + I(BA^2)) * (VH + LS + LMA + LCC)
+ I(VH^2) + I(LS^2 + I(LMA^2) + I(LCC^2))
+ (1 + VH + LS + LMA + LCC | site)
+ (1 + MAP + MAT + TSD + %Sand + %N + BA | sp)
+ offset(log(quadrats))
```

#### *Comparison between GLMM and CWMr approaches*

Test statistics in both approaches are calculated in the same way, as the ratio between the estimate of the corresponding parameter and its standard error. In the case of CWMr, test statistics are assumed to be *t*-distributed (i.e. p-values calculated from the Student's *t* distribution, here with 187 degrees of freedom) whereas they are assumed to be normally distributed in the case of GLMM.

**Table S4**. Possible outcomes and interpretations when comparing the agreement between the CWMr and GLMM approaches. Adapted from, or inspired by Laughlin et al. (2018).

<table>
<tr><td colspan="2" rowspan="2"></td><td colspan="2">GLMM</td></tr>
<tr><td>Non-significant</td><td>Significant</td></tr>
<tr><td rowspan="2">CWMr</td><td>Non-significant</td><td>Outcome 1:<br>The trait has no adaptive value along the environmental gradient.</td><td>Outcome 2:<br>The trait likely has adaptive value along the gradient, but the CWM values are being driven by other abiotic or biotic factors that vary spatially or temporally. These factors may blurry the trait-environment relationship at hand.</td></tr>
<tr><td>Significant</td><td>Outcome 3:<br><br>a) The relationship could be spurious since CWMr typically has inflated type I (Miller et al. 2019)<br><br>b) The relationship is not spurious, but the CWM values are not directly affected by the environmental variable in question. Other variables (possibly included in the GLMM) may be better explaining the variation of CWM values.</td><td>Outcome 4 – same sign:<br>The trait has adaptive value along the environmental gradient.<br><br>Outcome 5 – different sign:<br>Complete incongruence between methods, a mixture of outputs 2 and 3. The GLMM relationship would be closer to reflect the adaptive value of the trait, while that estimated by CWMr, if not spurious, would reflect the cumulative effects of other environmental factors.</td></tr>
</table>

### Appendix S2 – Supplementary Results

#### *Full GLMM results*

Overall, both temperature seasonality (TSD) and tree basal area (BA) had significant linear relationships with abundance (positive for TSD and negative for BA, as expected). Mean annual precipitation (MAP), mean annual temperature (MAT), and soil nitrogen content (%N) had unimodal, n-shaped relationships with abundance (negative quadratic terms significant at $p<0.01$). Soil sand content (%Sand) showed a similar unimodal relationship with abundance but with lower significance ($p<0.05$, Table S5). High values of %Sand and %N are associated with higher abundances due to significant positive linear terms (Table S5). Leaf size (LS) also had an overall positive linear effect on abundance. Leaf mass per area (LMA) had a negative quadratic relationship with abundance, and the optimal LMA value occurred at low LMA values (reflecting the negative linear term). Interactions between traits and environmental variables are described in the main text.

**Table S5.** Summary results of the generalized linear model (GLMM). Significance of model terms was evaluated using Type II Wald chi-square tests, i.e. according to the principle of marginality, each term is tested after all others in the same order or hierarchy, but ignoring the term's higher-order relatives (Fox and Weisberg 2019). In grey are shown the five most significant interactions, which are plotted in Fig. 3 in the main text. The marginal $R^2$ = 0.385, while the conditional $R^2$ was 0.977.

| Term | Estimate | Std. Error | Chisq | Pr(>Chisq) | |
|---|---|---|---|---|---|
| MAP | 0.026 | 0.086 | 0.218 | 0.6404 | |
| MAT | 0.045 | 0.176 | 1.142 | 0.2853 | |
| TSD | 0.405 | 0.101 | 12.989 | 0.0003 | *** |
| %Sand | 0.376 | 0.101 | 9.012 | 0.0027 | ** |
| %N | 0.526 | 0.107 | 23.715 | <0.0001 | *** |
| BA | -0.267 | 0.078 | 10.724 | 0.0011 | ** |
| MAP^2 | -0.202 | 0.058 | 17.403 | <0.0001 | *** |
| MAT^2 | -0.834 | 0.096 | 89.291 | <0.0001 | *** |
| TSD^2 | -0.140 | 0.061 | 1.951 | 0.1625 | |
| %Sand^2 | -0.145 | 0.081 | 4.489 | 0.0341 | * |
| %N^2 | -0.167 | 0.045 | 19.995 | <0.0001 | *** |
| BA^2 | -0.048 | 0.051 | 2.625 | 0.1052 | |
| VH | -0.219 | 0.209 | 2.129 | 0.1446 | |
| LS | 0.454 | 0.203 | 13.825 | 0.0002 | *** |
| LMA | -1.192 | 0.203 | 35.339 | <0.0001 | *** |
| LCC | 0.113 | 0.181 | 0.937 | 0.3332 | |
| VH^2 | -0.172 | 0.135 | 1.608 | 0.2048 | |
| LS^2 | -0.037 | 0.089 | 0.173 | 0.6776 | |
| LMA^2 | -0.323 | 0.115 | 7.860 | 0.0051 | ** |
| VH–MAP | 0.007 | 0.059 | 0.014 | 0.9066 | |
| LS–MAP | 0.011 | 0.057 | 0.039 | 0.8435 | |
| LMA–MAP | -0.053 | 0.058 | 0.839 | 0.3597 | |
| LCC–MAP | -0.030 | 0.044 | 0.457 | 0.4992 | |
| VH–MAT | 1.031 | 0.146 | 49.928 | <0.0001 | *** |
| LS–MAT | -0.067 | 0.144 | 0.218 | 0.6406 | |
| LMA–MAT | -0.332 | 0.144 | 5.314 | 0.0212 | * |
| LCC–MAT | -0.306 | 0.125 | 6.006 | 0.0143 | * |
| VH–TSD | 0.328 | 0.082 | 15.872 | 0.0001 | *** |
| LS–TSD | -0.242 | 0.082 | 8.764 | 0.0031 | ** |
| LMA–TSD | -0.239 | 0.082 | 8.583 | 0.0034 | ** |
| LCC–TSD | 0.131 | 0.070 | 3.456 | 0.0630 | . |
| VH–%Sand | 0.047 | 0.077 | 0.379 | 0.5381 | |
| LS–%Sand | -0.266 | 0.075 | 12.634 | 0.0004 | *** |
| LMA–%Sand | 0.096 | 0.076 | 1.616 | 0.2037 | |
| LCC–%Sand | 0.135 | 0.062 | 4.705 | 0.0301 | * |
| VH–%N | 0.172 | 0.086 | 4.000 | 0.0455 | * |
| LS–%N | -0.088 | 0.084 | 1.089 | 0.2967 | |
| LMA–%N | -0.200 | 0.086 | 5.403 | 0.0201 | * |
| LCC–%N | -0.070 | 0.073 | 0.935 | 0.3337 | |
| VH–BA | -0.035 | 0.057 | 0.365 | 0.5459 | |
| LS–BA | -0.074 | 0.055 | 1.814 | 0.1780 | |
| LMA–BA | -0.085 | 0.057 | 2.279 | 0.1311 | |
| LCC–BA | 0.104 | 0.045 | 5.318 | 0.0211 | * |

**Table S5**. Continued.

| Term | Estimate | Std. Error | Chisq | Pr(>Chisq) | |
|---|---|---|---|---|---|
| VH–MAP | 0.055 | 0.038 | 2.067 | 0.1505 | |
| LS–MAP | 0.011 | 0.035 | 0.090 | 0.7644 | |
| LMA–MAP | -0.040 | 0.036 | 1.221 | 0.2692 | |
| LCC–MAP | 0.000 | 0.025 | 0.000 | 0.9973 | |
| VH–MAT | -0.060 | 0.063 | 0.903 | 0.3421 | |
| LS–MAT | 0.106 | 0.059 | 3.174 | 0.0748 | . |
| LMA–MAT | 0.086 | 0.061 | 1.970 | 0.1605 | |
| LCC–MAT | 0.067 | 0.043 | 2.415 | 0.1202 | |
| VH–TSD | -0.086 | 0.039 | 4.839 | 0.0278 | * |
| LS–TSD | 0.139 | 0.037 | 13.853 | 0.0002 | *** |
| LMA–TSD | 0.123 | 0.037 | 10.903 | 0.0010 | *** |
| LCC–TSD | 0.008 | 0.025 | 0.103 | 0.7481 | |
| VH–%Sand | -0.052 | 0.052 | 1.019 | 0.3128 | |
| LS–%Sand | 0.004 | 0.049 | 0.007 | 0.9352 | |
| LMA–%Sand | 0.033 | 0.049 | 0.452 | 0.5015 | |
| LCC–%Sand | -0.035 | 0.033 | 1.156 | 0.2823 | |
| VH–%N | 0.009 | 0.030 | 0.093 | 0.7600 | |
| LS–%N | -0.015 | 0.028 | 0.300 | 0.5841 | |
| LMA–%N | 0.031 | 0.029 | 1.129 | 0.2881 | |
| LCC–%N | 0.040 | 0.020 | 4.041 | 0.0444 | * |
| VH–BA | 0.033 | 0.033 | 0.981 | 0.3221 | |
| LS–BA | 0.018 | 0.030 | 0.335 | 0.5628 | |
| LMA–BA | 0.007 | 0.031 | 0.051 | 0.8220 | |
| LCC–BA | -0.023 | 0.022 | 1.137 | 0.2862 | |

**Table S6.** Akaike information criterion (AIC) for models nested within the full model detailed in Table S5. Nested models result from removing either of both interaction and quadratic terms from the full model. ΔAIC measures the difference (increase) in AIC between the full model and the corresponding nested model. Following equation 1 in the main text, a generic equation of the fixed part of the model corresponding to each trait-environment is shown below each model's name. The random part is the same in all models. The number of parameters estimated by each model is also shown.

| Model | Terms removed from the full model | AIC | ΔAIC | Parameters |
|---|---|---|---|---|
| Linear terms for T and E<br>$\beta_0 + \beta_1 T_i + \beta_3 E_j$ | Quadratic and interaction terms | 57115 | 124 | 55 |
| Linear and quadratic terms for T and E<br>$\beta_0 + \beta_1 T_i + \beta_2 T_i^2 + \beta_3 E_j + \beta_4 E_i^2$ | Interaction terms | 57041 | 50 | 61 |
| Linear and interaction terms for T and E<br>$\beta_0 + \beta_1 T_i + \beta_3 E_j + \beta_5 T_i E_j$ | Quadratic terms | 57076 | 85 | 79 |
| Full model<br>$\beta_0 + \beta_1 T_i + \beta_2 T_i^2 + \beta_3 E_j$<br>$+\beta_4 E_i^2 + \beta_5 T_i E_j + \beta_6 T_i E_j^2$ | None | 56991 | 0 | 113 |

#### *Results of CWM regressions*

Here we expand the results shown in Fig. 4 in the main text. We show the actual values of the *t* statistics associated with linear and quadratic terms (Table S7). The table also shows a qualitative description of the shape of each significant trait–environment relationship according to this approach.

**Table S7.** Summary of trait–environment (T–E) relationships estimated by the community-weighted mean (CWM) regression approach. Significant positive relationships are highlighted in green and negative in orange (critical *t* for 187 degrees of freedom is 2.602). Alternative relationship types are exclusively rectilinear or directional (only $\gamma_1 \neq 0$), curved with a directional component (both $\gamma_1 \neq 0$, $\gamma_2 \neq 0$), and exclusively curved, with no significant directionality (only $\gamma_2 \neq 0$).

| **T–E** | **Estimate** | **SE** | ***t* value** | **Pr(>\|t\|)** | **Estimate** | **SE** | ***t* value** | **Pr(>\|t\|)** | **Type** |
|---|---|---|---|---|---|---|---|---|---|
| LMA–MAT | -0.285 | 0.030 | -9.526 | <0.00001 | 0.027 | 0.025 | 1.113 | 0.26702 | Directional |
| VH–%Sand | -0.182 | 0.027 | -6.845 | <0.00001 | -0.069 | 0.028 | -2.408 | 0.01702 | Directional |
| LCC–MAT | -0.155 | 0.033 | -4.699 | 0.00001 | 0.065 | 0.027 | 2.390 | 0.01783 | Directional |
| LMA–MAP | -0.128 | 0.028 | -4.527 | 0.00001 | -0.032 | 0.022 | -1.450 | 0.14872 | Directional |
| LS–%Sand | -0.116 | 0.021 | -5.420 | <0.00001 | -0.054 | 0.023 | -2.367 | 0.01898 | Directional |
| LS–MAP | 0.060 | 0.023 | 2.619 | 0.00956 | 0.013 | 0.018 | 0.713 | 0.47666 | Directional |
| VH–MAP | 0.116 | 0.029 | 4.026 | 0.00008 | 0.038 | 0.023 | 1.680 | 0.09464 | Directional |
| LS–MAT | 0.117 | 0.034 | 3.406 | 0.00081 | 0.013 | 0.028 | 0.469 | 0.63960 | Directional |
| LCC–%Sand | 0.138 | 0.024 | 5.665 | <0.00001 | 0.035 | 0.026 | 1.338 | 0.18237 | Directional |
| VH–nitro | 0.214 | 0.026 | 8.323 | <0.00001 | -0.024 | 0.015 | -1.620 | 0.10689 | Directional |
| LMA–%Sand | 0.233 | 0.024 | 9.736 | <0.00001 | 0.061 | 0.026 | 2.369 | 0.01887 | Directional |
| VH–MAT | 0.275 | 0.031 | 8.774 | <0.00001 | -0.035 | 0.026 | -1.376 | 0.17063 | Directional |
| LMA–nitro | -0.208 | 0.025 | -8.389 | <0.00001 | 0.058 | 0.015 | 3.960 | 0.00011 | Curved & directional |
| LCC–nitro | -0.162 | 0.023 | -7.044 | <0.00001 | 0.053 | 0.013 | 3.963 | 0.00011 | Curved & directional |
| LS–TSD | -0.078 | 0.022 | -3.546 | 0.00049 | 0.061 | 0.020 | 3.117 | 0.00211 | Curved & directional |
| LS–nitro | 0.060 | 0.022 | 2.713 | 0.00730 | -0.052 | 0.013 | -3.988 | 0.00010 | Curved & directional |
| LCC–MAP | -0.057 | 0.026 | -2.211 | 0.02824 | -0.054 | 0.020 | -2.687 | 0.00785 | Curved |
| LMA–BA | -0.056 | 0.030 | -1.892 | 0.06005 | 0.037 | 0.023 | 1.593 | 0.11292 | NS |
| LMA–TSD | 0.075 | 0.029 | 2.584 | 0.01054 | -0.032 | 0.026 | -1.229 | 0.22061 | NS |
| VH–TSD | -0.031 | 0.030 | -1.042 | 0.29858 | 0.031 | 0.027 | 1.150 | 0.25182 | NS |
| LCC–BA | 0.002 | 0.027 | 0.080 | 0.93669 | 0.023 | 0.021 | 1.090 | 0.27724 | NS |
| LCC–TSD | 0.051 | 0.026 | 1.924 | 0.05590 | -0.008 | 0.023 | -0.320 | 0.74937 | NS |
| VH–BA | 0.035 | 0.030 | 1.137 | 0.25686 | 0.005 | 0.024 | 0.210 | 0.83403 | NS |
| LS–BA | -0.041 | 0.023 | -1.764 | 0.07932 | -0.002 | 0.018 | -0.113 | 0.90994 | NS |
